# Cilia-mediated cerebrospinal fluid flow modulates neuronal and astroglial activity in the zebrafish larval brain

**DOI:** 10.1101/2024.02.01.578354

**Authors:** Percival P. D’Gama, Inyoung Jeong, Andreas Moe Nygård, Ahmed Jamali, Emre Yaksi, Nathalie Jurisch-Yaksi

## Abstract

The brain uses a specialized system to transport cerebrospinal fluid (CSF). This system consists of interconnected ventricles lined by ependymal cells, which generate a directional flow upon beating of their motile cilia. Motile cilia act jointly with other physiological factors, including active CSF secretion and cardiac pressure gradients, to regulate CSF dynamics. The content and movement of CSF are thought to be important for brain physiology. Yet, the link between cilia-mediated CSF flow and brain function is poorly understood. In this study, we addressed the role of motile cilia-mediated CSF flow on brain development and physiology using zebrafish larvae as a model system. By analyzing mutant animals with paralyzed cilia, we identified that loss of ciliary motility did not alter progenitor proliferation, overall brain morphology, or spontaneous neural activity. Instead, we identified that cilia paralysis led to randomization of brain asymmetry. We also observed altered neuronal responses to photic stimulation, especially in the optic tectum and hindbrain. Since astroglia contact CSF at the ventricular walls and are essential for regulating neuronal activity, we next investigated astroglial activity in motile cilia mutants. Our analyses revealed a striking reduction in astroglial calcium signals both during spontaneous and light-evoked activity. Altogether, our findings highlight a novel role of motile cilia-mediated flow in regulating brain physiology through modulation of neural and astroglial networks.

## Introduction

Motile cilia are microtubule-based organelles that are located at the surface of specific cells (1) and actively beat to generate motion or pump fluids. In the brain, motile cilia are found on ependymal cells, which are specialized cell types lining the brain ventricles (2-8). Ciliary beating contributes to cerebrospinal fluid (CSF) circulation together with CSF secretion, pressure gradients related to the cardiac cycle and respiration, and bodily movement (4, 9, 10). Collectively these mechanisms ensure CSF flows across the brain and spinal cord (4, 5, 11-14), which is necessary for brain homeostasis and waste removal (11).

Despite clear evidence showing that ciliary beating generates CSF movement along the ventricular wall (4, 8, 12, 13, 15, 16), the function of motile cilia-mediated flow in the brain remains highly debated. Motile cilia have been causally linked to hydrocephalus, a condition with enlarged ventricles, especially in rodents (5, 17-19), but the association between hydrocephalus and motile cilia defects remains weak in humans (17, 20-22). It has been postulated that the difference in brain anatomy and physiology may account for the increased prevalence of hydrocephalus in rodents as compared to human (17). Since cilia mainly contribute to the near-wall and not the bulk CSF flow, one could predict that cilia will have less impact when ventricles are big, such as in humans, and thereby a minor causal role in hydrocephalus. Nevertheless, motile cilia will generate a directional fluid flow at the surface of the ventricle and as such have been suggested to contribute to the establishment of signaling molecule gradients within the brain (23, 24). This was experimentally shown by one study so far, which has observed abnormal neuronal migration during postnatal brain development in a cilia mutant animal (13). Due to the high prevalence of hydrocephalus in motile cilia mutant mice (20, 25), it has remained difficult to assess the function of motile cilia on brain physiology *per se*, independently of ventricular defects or altered intercranial pressure.

In our study, we leveraged the small size and genetic amenability of zebrafish to specifically address these questions. We used a motile cilia mutant, *schmalhans* (*smh*, carrying a mutation in *ccdc103* gene) (26) that lack motile cilia-mediated flow in the zebrafish larval brain, without displaying altered ventricular size (4). We found that the absence of motile cilia-mediated flow in larval zebrafish had no effects on brain morphology, progenitor cell proliferation or spontaneous brain activity. Instead, we found that loss of ciliary motility led to altered light-evoked neuronal activity and brain asymmetry. Furthermore, we observed reduced spontaneous and sensory-driven astroglial calcium signals. Overall, our findings indicate that loss of motile cilia-generated flow affects brain physiology in larval zebrafish, most likely through altered astroglial function. Altogether, our findings shed new light on the complex interactions between motile cilia function, neuronal responses, and astroglial networks in the context of brain development and physiology.

## Results

### Loss of ciliary motility does not impair the overall brain anatomy, the proliferation of progenitors or the number of microglia in larval zebrafish

To investigate the function of motile cilia-mediated flow in the developing brain, we focused our analyses on a zebrafish motile cilia mutant line, *ccdc103*^*tn222a*^ (referred to as *smh*), which display immotile cilia due to a disrupted assembly of the dynein arms (26). In agreement with our prior work (4, 8), we observed that *smh* mutants assemble glutamylated-positive cilia at 4 days post-fertilization (dpf) in all motile cilia lineages of the brain like controls **(figure 1A1-B4)**. However, since cilia are immotile, they do not generate fluid flow (4), thereby allowing us to specifically study the impact of ciliary motility rather than the loss of motile cilia in the developing brain.

**Figure 1:**
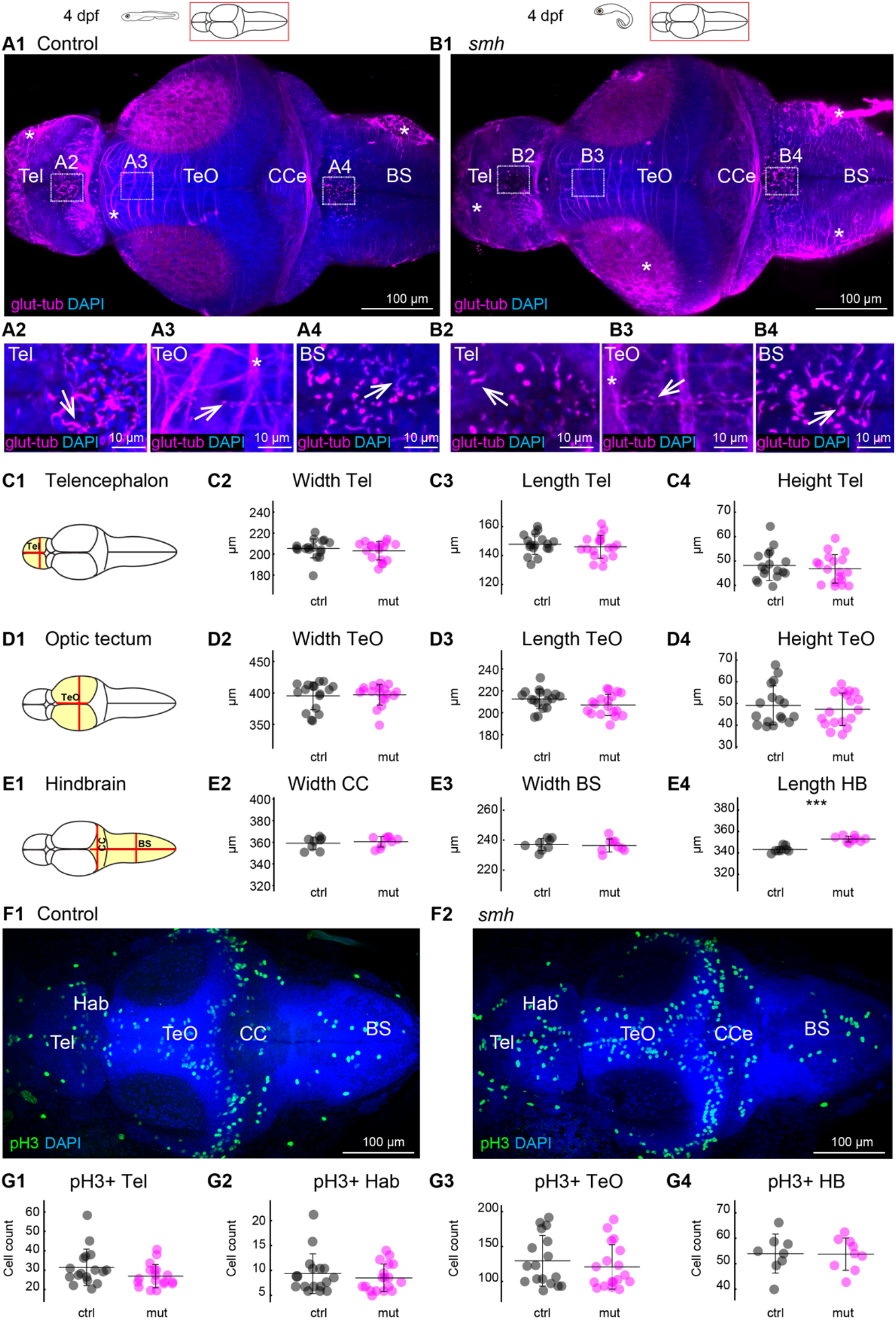
Loss of cilia motility does not impact the overall brain morphology and progenitor proliferation. (**A1-A4** and **B1-B4)** Staining of dissected 4 dpf brains with glutamylated tubulin used to identify motile cilia, n=4 controls and mutants. **(A1)** At 4 dpf, single glutamylated tubulin-positive cilia were present in the forebrain choroid plexus **(A2)**, on the dorsal roof and ventral part **(A3)** of the tectal/diencephalic ventricle and in the rhombencephalic choroid plexus in the brain stem **(A4). (B1)** In smh mutant, glutamylated tubulin-positive cilia are still present in the forebrain choroid plexus **(B2)**, optic tectum **(B3)**, and the rhombencephalic choroid plexus in the brain stem **(B4). (C1-E4)** Quantification of brain morphology of 4 dpf larval brains for control (black) and smh mutants (magenta). Schematic representation of the measurements for **(C1)** telencephalon, **(D1)** optic tectum, **(E1)** hindbrain regions. Brain size was estimated by measuring the width, length, and height of telencephalon **(C2, C3, C4)**, optic tectum **(D2, D3, D4)**, width of CCe **(E2)**, width of BS **(E3)**, length of hindbrain **(E4)**. Controls are in black and smh mutants in magenta. n =18 controls and 19 mutants for **C2, C3, C4, D2, D3, D4, D2, D4**. n = 9 controls and 10 mutants for **E2, E3** and **E4. (F1, F2)** staining for mitotic cells using an anti-pH3 antibody. **(G1-G4)** Cell count for pH3 positive cells (pH3+) in telencephalon **(G1)**, habenula **(G2)**, optic tectum **(G3)** and hindbrain **(G4)**. ^***^: p < 0.001 according to Wilcoxon Rank Sum test. Mean +/-standard deviation is indicated on scatter plots. Tel, Telencephalon; TeO, Optic Tectum; BS, Brain stem; CCe, Corpus Cerebelli; HB, Hindbrain, Hab: habenula.

We performed various measurements of brain size in the motile cilia mutants, including the width, length, and height of the telencephalon **(figure 1C1-C4)**, optic tectum **(figure 1D1-D4)**, and hindbrain **(figure 1E1-E4)**. We did not find significant differences in any measurements **(figure 1C1-E4)**, except the length of the hindbrain **(figure 1E4)**, which may relate to the body curvature of the mutants. Additionally, our staining with glutamylated tubulin, which labels axons and cilia, did not reveal any gross abnormality in brain patterning **(figure 1A-B)**.

Neural progenitor cells are located at the ventricular zone in direct contact with CSF (27-29). Since their proliferation can be regulated by the molecular content and dynamics of CSF (30-32), we investigated the impact of disrupted cilia-mediated flow on cell proliferation. To this end, we stained mitotic cells using a phosphorylated Histone H3 (pH3) antibody **(figure 1F1-F2)**. Our analyses showed no significant differences in the number and location of dividing cells across brain regions **(figure 1G1-G4)**.

Previous work in zebrafish suggested that loss of CSF flow leads to the activation of immune cells and eventually scoliosis (33). To assess if loss of ciliary motility results in an increased number of immune cells in the brain at larval stage, we stained microglia cells **(figure S1A1, A2)** using the 4C4 antibody (34). Our analysis revealed no significant differences in the number and location of microglia in all brain regions **(figure S1B1-B4)**. Taken together, our findings indicate that the absence of cilia-driven flow has minor effects on brain morphology and does not affect the number and location of dividing cells or microglia in the larval brain.

### Transcriptomic analysis identifies differentially regulated genes involved in ciliary function and movement

To explore the impact of loss of ciliary motility on gene expression, we performed RNA-sequencing and transcriptomic analysis of whole 4 dpf larvae. We identified a total of 42 differentially expressed genes (DEGs), comprising 36 upregulated DEGs and 6 downregulated DEGs **(figure 3A-B)**, between control and *smh* mutant larvae. We classified the genes into different Gene Ontology (GO) terms consisting of GO biological process, GO cellular component, and GO molecular function **(figure 2C** and **S2A-C)**. Remarkably, the prominent GO categories involved processes related to ciliary assembly, ciliary movement, and cytoskeleton. Notably, genes related to ciliary motility, such as dynein arms genes *dnah2, dnah5l, dnah6, dnah7* **(figure 2B1)** and the master transcriptional regulator of motile ciliogenesis *foxj1a* **(figure 2B2)** were upregulated in the motile cilia mutant. The CSF-contacting neurons’ expressing gene *urotensin-related peptide 2* (*urp2*) was downregulated in agreement with its described function in the curved body axis of cilia mutants (35). No GO terms associated with brain development or signaling were statistically significant. Taken together, our results show that loss of cilia-mediated flow mainly impacts the expression of genes associated with ciliary motility.

**Figure 2:**
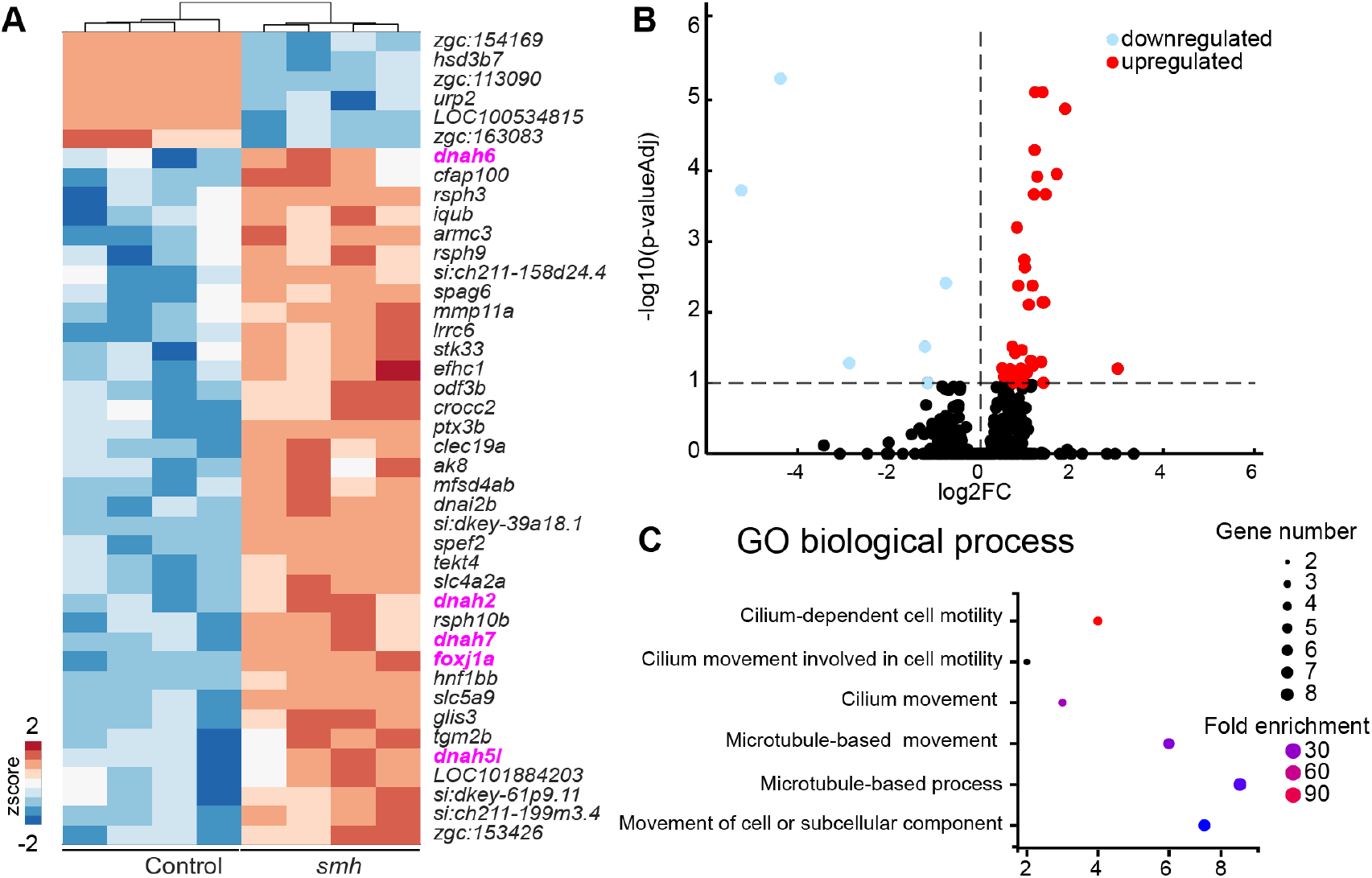
RNA sequencing identifies differentially regulated genes (DEG) involved in ciliary movement and dynein activity. **(A)** Heatmap of expression of DEGs from control and smh 4 dpf whole larvae (n=4 RNA preparation obtained from circa 30 control and smh) shown as z-score. Increased expression is represented with red and decreased expression with blue. A total of 42 (36 upregulated and 6 downregulated) differentially expressed mRNAs were identified. **(B)** Volcano plot showing the DEGs in smh as compared with control. The horizontal dotted line indicates the P adjusted value of 0.1. The vertical dotted line separate upregulated (red) and downregulated (blue) DEGs. **(C)** List of Gene Ontology (GO) biological process that are significantly enriched. See supplementary table 1.

**Figure 3:**
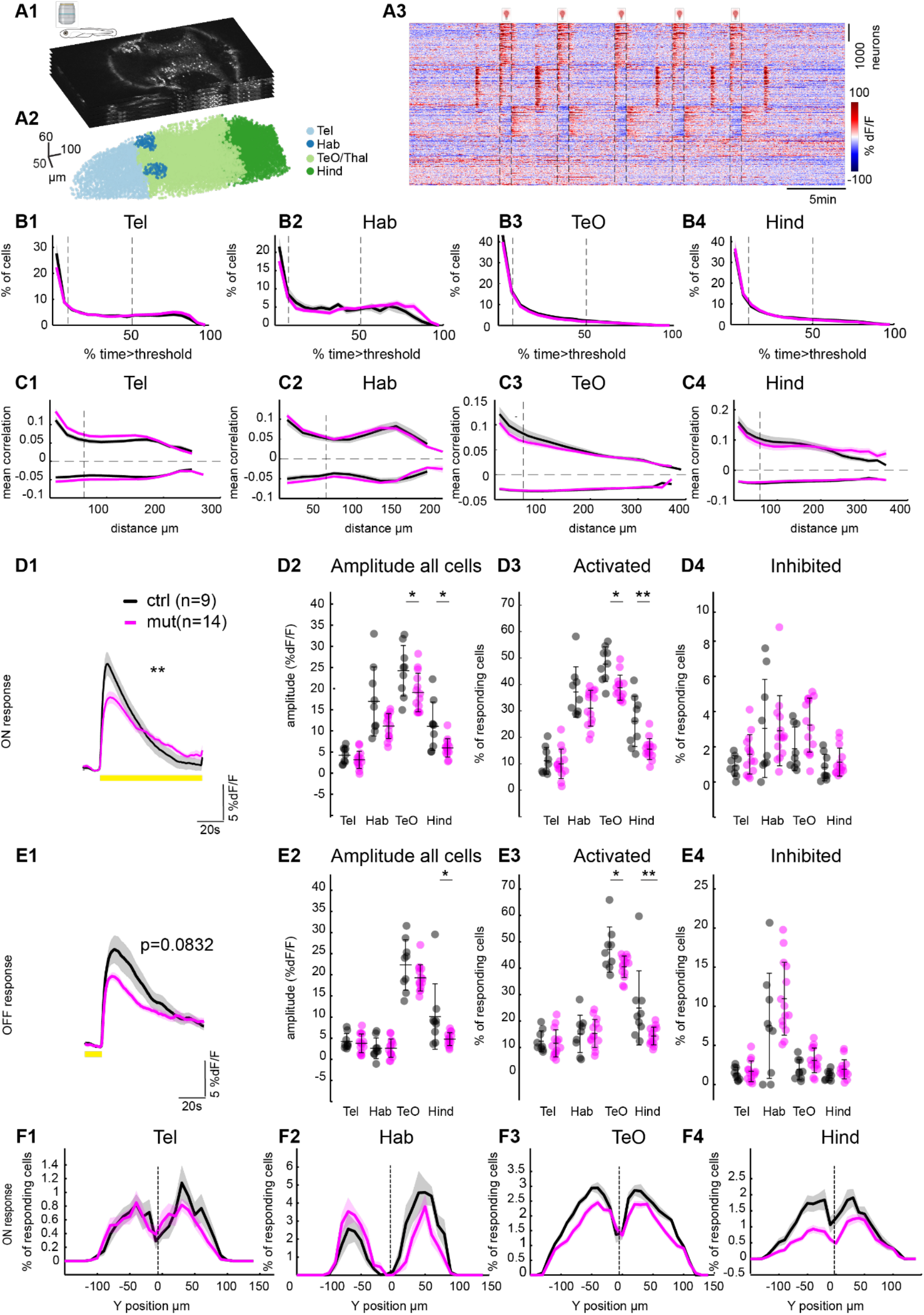
Impaired neuronal activity and brain asymmetry in motile cilia mutants. **(A1)** Optical sections of multiplane recording of transgenic zebrafish larvae expressing nuclear GCaMP6s in all neurons Tg(elavl3:H2BGcaMP6s). **(A2)** Color coded segmented nuclei from different brain regions identified by anatomical landmarks. **(A3)** Neuronal activity represented as change of fluorescence (% dF/F) in one representative control. Traces were sorted based on their activity using k-means clustering (warm color represents higher calcium signals). **(B1-B4)** Percentage of activity of neurons in Telencephalon **(B1)**, Habenula **(B2)**, TeO/thalamus **(B3)** and Hindbrain **(B4)**, for control (black) and smh (magenta). (+/-standard error of mean as the shaded area). Dotted lines indicate cells that are active 10% and 50% of the time period. **(C1-C4)** Mean Pearson’s correlation versus distance showing positive and negative correlations between cells in Telencephalon **(C1)**, Habenula **(C2)**, TeO/thalamus **(C3)** and Hindbrain **(C4)**, for control (black) and mutant (magenta). Significance tests for correlation were computed for cells located within 60 µm indicated by the dotted line on the X-axis. **D1, E1** Average response of neuronal activity to light stimulation for control (black) and mutant (magenta) for ON **(D1)** and OFF **(E1)** response conditions (+/-standard error of mean as the shaded area). **D2, E2** Average amplitude of all cells during ON **(D2)** and OFF **(D3)** response for control (black) and mutant (magenta), amplitude was significantly reduced in TeO and Hind regions for ON response and only in hindbrain brain regions for OFF response. **D3, E3** and **D4, E4** Percentage of cells that are activated during ON **(D3)** and OFF response **(E3)** and inhibited during ON **(D4)** and OFF response **(E4). (F1-F4)** Y position for responding cells in the ON response for telencephalon **(F1)**, Habenula **(F2)**, optic tectum **(F3)**, and hindbrain **(F4)**. Dotted line at 0 represents the position of the midline where the ventricles are located ^*^: p <0.05, ^**^: p <0.01 according to Wilcoxon Rank Sum test. Mean+/-standard deviation is indicated on scatter plots. n = 9 controls and 14 mutants, control (black) and smh mutant (magenta). Tel, Telencephalon; TeO, Optic Tectum; Hind, Hindbrain, Hab: Habenula.

### Altered light-evoked neural activity and loss of brain asymmetry in motile cilia mutant

To identify whether cilia-mediated flow modulates brain physiology, we measured spontaneous and sensory-driven neural activity in 4 dpf *smh* larvae by two-photon calcium imaging, when mutant larvae remain relatively healthy beside their curved body axis. We performed a 40 min long recording consisting of 10 min of spontaneous activity, followed by a series of photic stimulations that measure visual responses and can trigger hyperexcitability (36). More specifically, we recorded 8 volumetric planes of transgenic larvae expressing the nuclear calcium indicator GCaMP6s in all neurons *(Tg(elavl3:H2B-GCaMP6s))* **(figure 3A1)** (37) and collected data from four brain areas: the telencephalon, optic tectum/thalamus, habenula, and hindbrain **(figure 3A2)**. As shown for a representative example **(figure 3A3)**, we observed distinct clusters of neurons exhibiting varying levels of activity during photic stimulations.

First, we assessed whether loss of ciliary motility influenced spontaneous brain activity by analyzing neural activity during the period preceding the light stimuli in control and *smh* mutant groups. To this end, we measured the activity of cells by quantifying the percentage of time they spent above a threshold of four times the baseline and generated an average histogram for each brain regions. Our analysis showed no significant differences between control and mutant groups in the activity of cells in all brain regions **(figure 3B1-B4** and **figure S3A1-D2)**. Furthermore, we investigated whether cells were differently synchronized with each other. To this end we calculated the mean Pearson’s correlations as a function of the distance between pairs of cells within distinct brain regions (38) Our analysis revealed no significant differences of neural synchrony in *smh* mutants as compared to controls **(figure 3C1-C4** and **figure S3E1-H2)**.

Next, we analyzed neuronal activity during the photic stimulation. We first quantified the average activity of all neurons and found a significantly reduced response for the ON **(figure 3D1)** but not the OFF **(figure 3E1)** response. By measuring the response amplitude for each brain region separately **(figure 3D2** and **E2)**, we identified reduced activity in the visual center, known as the optic tectum, and hindbrain for the ON response **(figure 3D2)** and only in the hindbrain for the OFF response **(figure 3E2)**. We then investigated whether cells were generally less active or whether there were fewer responding cells. To determine the ON and OFF responding cells, we analyzed 10 seconds after the stimulus and categorized cells with a mean amplitude greater than 2 standard deviations as responding. Our findings revealed a significant reduction in the number of activated neurons in the optic tectum and hindbrain for both the light ON **(figure 3D3)** and OFF conditions **(figure 3E3)**. However, there were no significant differences in the number of inhibited neurons for either light ON **(figure 3D4)** or OFF **(figure 3E4)** conditions. Altogether, by showing altered photic responses in motile cilia mutants, our results suggest that the flow generated by motile cilia is necessary to maintain normal brain excitability.

The retina serves as the first point of light detection and retinal defects could lead to reduced photic activity. Therefore, we analyzed the retinal phenotype of motile cilia even though motile cilia have not been described in the retina. To this end, we first prepared retinal cryosections of 4 dpf control and *smh* larvae **(figure S4A1, B1)** and stained the photoreceptor outer segments using the lipophilic stain Dil. We observed normal intact elongated outer segment in both controls **(figure S4A2)** and *smh* mutants **(figure S4B2)**. Next, we performed DiI-based tracing of retinal ganglion cells axons to the optic tectum to identify potential mis-patterning that could explain the reduced photic activity in *smh* mutants. Our results did not indicate any projection defects **(figure S4C1, C2)**. Finally, we investigated retinal function by electroretinography (ERG) **(figure S4D1, D2)**, using a train of 1s-long blue light stimuli. In agreement with intact photoreceptors’ outer segments, we observed normal ERG signals in the 4dpf *smh* mutants as compared to controls **(figure S4D1, D2)**. Taken together, our analyses suggest that reduced photic activity does not stem from retinal dysfunction or mis-patterning of retinal ganglion cells axons but rather due to reduced activation of neurons in the optic tectum.

Given that the motile ciliated cells line the ventricles of the zebrafish brain (4, 8), we were interested to see if the reduced neuronal responses were limited to the cells lining the ventricles. To investigate this, we identified the location of responding cells in relation to the midline of the brain where the ventricles are located. More specifically, we reported the percentage of responding cells along the medial-lateral axis (y axis) of our recordings, with a medial position (y value of 0) being closer to the ventricle and lateral position (y value <50 or <-50) being further away from the ventricle. Our observations revealed that the diminished neuronal responses in the optic tectum and hindbrain are visible across the width of the brain regions and do not depend on the medio-lateral position of the neurons **(figure 3F1-F4** and **S5A-D)**.

Interestingly, responses of the habenula, which are normally asymmetrical following light stimulation (39, 40), were randomized in the *smh* mutant **(figure 3F2, S6A)**. This finding suggests a disrupted establishment of asymmetry in the habenula upon motile cilia defects. To gain deeper insights into this phenomenon, we analyzed axonal projections to the habenula. We focused our analysis on olfactory bulb neurons that project solely to the right habenula using the *Tg(lhx2a:gap-YFP)* transgenic zebrafish line (41). In agreement with our functional data, we observed a remarkable disparity between the control and *smh* mutant groups. While in the control zebrafish, axons projected solely to the right habenula as expected **(figure S6B1)** (41, 42), there was a prominent prevalence of axonal projections to the left habenula in *smh* mutants **(figure S6B2)**. Some mutants even exhibited a complete absence of axonal projection from the olfactory bulb to the habenula **(figure S6B3**-**4)**. Interestingly, we did not observe asymmetry defects in the habenula of *elipsa* mutants (**figure S7A**,**B)**, which loose cilia between 10 somites and 30 hpf, before motile cilia are formed in the brain (4), but after left-right asymmetry is established (43). Given the well-established role of motile cilia in the left-right asymmetry breaking organ, known as the Kupffer’s vesicle (44-46), our results suggest that habenular asymmetry may be driven by the Kupffer’s vesicle during development.

Altogether, our findings indicate that motile cilia mutants exhibit defects in brain asymmetry and reduced photic activity in the optic tectum and hindbrain.

### Reduced spontaneous and sensory-driven astroglia activity in the *smh* mutants

We next sought to study the impact of cilia-mediated flow on astroglia cells since their soma are in direct contact with CSF. Moreover, astroglia play critical roles in supporting and regulating neuronal function, including maintaining ion balance, providing structural support, and modulating neurotransmitter levels (29, 47, 48). To measure astroglia activity, we performed the same photic stimulation protocol as described above but used the zebrafish line *Tg(GFAP:gal4)Tg(UAS:Gcamp6s)* expressing the calcium indicator GcaMP6 in all astroglia cells (36, 48, 49). We calculated the change of fluorescence intensity in a region of interest encompassing the soma of all astroglial cells which are located at the midline and lining the ventricles **(figure 4A)**, as previously described (36, 48, 49). Our analysis revealed that on average astroglial cells were generally less active in *smh* mutants as compared to controls throughout the recording **(figure 4B1, B2** and **C)**. We also observed a significantly reduced response to the light ON stimulus in the mutant animals **(figure 4D, E)**. Altogether, our results highlight a role of motile cilia-mediated flow in regulating astroglial function, which in turn may explain the altered neuronal physiology.

**Figure 4:**
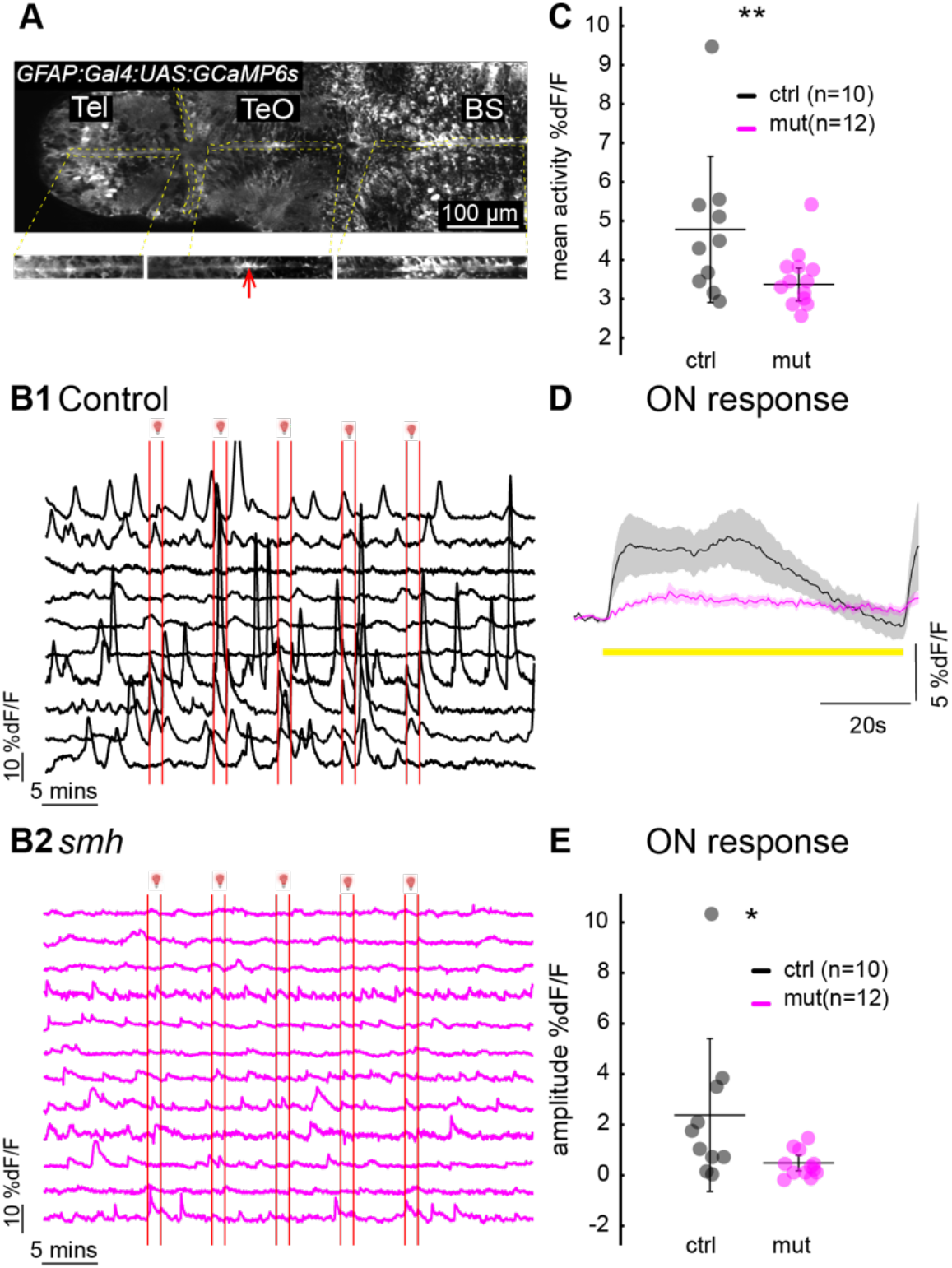
Reduced spontaneous and photic-induced astroglial activity in the smh larvae. **(A)** Optical section of multiplane recording of transgenic zebrafish larvae expressing GCaMP6s in glia cells Tg(GFAP:gal4)Tg(UAS:GCaMP6s). Regions of interest (ROI) were drawn at the soma of the cells lining the ventricles (red arrow drawn on inset) from each optical plane. **(B1-B2)** Averaged glia cell activity for the entire 40-minute recording in control **(B1)** and smh mutant **(B2)**. Each larva was subjected to 5 light flashes of 1 minute (light window indicated using red lines). **(C)** Mean glia activity for control (black) and smh mutant (magenta). **(D)** Average response of glia activity to light stimulation for control (black) and mutant (magenta) for ON response (+/-standard error of mean as the shaded region trace). **(E)** Amplitude of glial responses during ON response. ^*^: p <0.05, ^**^: p<0.01 according to Wilcoxon Rank Sum test. Mean+/-standard deviation is indicated on scatter plots. n = 10 controls and 12 mutants, control (black) and smh mutant (magenta). Tel, Telencephalon; TeO, Optic Tectum; BS, Brain stem.

## Discussion

In this study, we addressed the debated function of motile cilia in the brain, leveraging the genetic advantages of zebrafish. Unlike previous studies in rodents, our research utilized a mutant zebrafish model that lacked cilia-mediated flow but did not have altered ventricular sizes, thereby allowing us to study the impact of motile cilia independently from ventricular abnormalities. Using this mutant, we found that cilia-mediated flow does not impact brain morphology, cell proliferation and immune cells in larval stages. Instead, our study uncovered novel insights into the role of motile cilia in brain physiology. We observed significant alterations in light-evoked neuronal activity, brain asymmetry, and astroglial calcium levels in larval zebrafish lacking ciliary motility. These findings shed new light on the intricate relationship between motile cilia function, neuronal responses, and glial networks, offering a fresh perspective on the role of motile cilia in brain development and physiology.

For this study, we specifically selected a mutation affecting ciliary motility and not ciliogenesis to be able to disentangle the function of fluid flow from other potential signaling function of motile cilia. Our cilia staining showed indeed no observable changes in motile cilia formation. It is thereby consistent with the initial work from (26) reporting normal ciliogenesis, yet total paralysis of cilia in *smh* mutants in the pronephros, olfactory placode and in spinal canal, and our previous research in the larval *smh* brain (4).

Dysfunction of motile cilia in rodent models is very strongly associated with the development of an enlarged ventricular system, known as hydrocephalus (5, 17). We previously reported a normal sized ventricular system in the *smh* mutant at larval stages (4). Since hydrocephalus can lead to secondary brain damage (50), especially in cases of increased intracranial pressure, we argue that unlike mice which develops hydrocephalus upon motile cilia dysfunction, our zebrafish model is well-suited to study the direct effects of cilia-mediated flow on the brain. Notably, we analyzed the impact of cilia paralysis at 4 dpf, which is circa 2.5 days after the onset of ciliary beating (4).

Our analysis did not reveal a reduction in number of dividing cells in any brain regions in the motile cilia mutants, suggesting that cilia-mediated fluid flow does not impact cell proliferation in the brain. This is rather puzzling given the proximity of progenitors to the CSF and the fact that CSF carries important signaling molecules for cell proliferation (31, 51). One could have expected that CSF movement is involved in the delivery of signaling and pro-proliferative molecules in the brain. Besides chemical factors delivered by CSF, mechanical forces on brain tissues and cells were also shown to regulate the proliferation of progenitors (52). Notably, neuronal proliferation in the subventricular zone of the adult mouse brain is regulated by mechanical forces through the somatic epithelial sodium channel ENaC (30). Piezo1-dependent mechanosensing was also shown to regulate neuronal differentiation in stem cells (53) and neuronal proliferation in the dentate gyrus through the release of ATP by astrocytes (54). In our previous work (4), we identified that bodily movements induce a strong CSF displacement in the brain ventricles. This could explain why we do not see any differences in cell proliferation in the *smh* mutants, as other physiological factors, including bodily movement and heartbeat, will compensate for CSF movement and mechanical forces.

Our transcriptomic analysis of *smh* mutants revealed a surprisingly low number of differentially expressed genes. We mainly observed upregulation of genes associated with ciliary movement including the master regulator of motile ciliogenesis *foxj1a* and some of its target genes, especially *dnah* genes, which encode the dynein arms of motile cilia (55, 56). This indicates that dysfunctional motile cilia can lead to the activation of a transcriptional program trying to compensate ciliary paralysis. This agrees well with the described *foxj1a* upregulation in scoliotic juvenile zebrafish with *sco-spondin* mutations (57) and in response to epithelial stretch following injury in the developing kidney (58). Our analysis also revealed downregulation of *urotensin-related peptide* 2 (*urp2)*, which has been associated with the body curvature (35) and scoliosis (57) of cilia mutants. Earlier studies on scoliotic zebrafish mutants with underlying ciliary dysfunction revealed increased neuroinflammation at juvenile stage (33, 57). In our analyses, we did not detect any alteration in either microglia number or transcriptomics signature of inflammation in *smh* mutants at larval stage, suggesting that neuroinflammation may occur later in life, at juvenile stage, when the animals develop scoliosis. Our transcriptomic analysis also did not pick up differentially regulated genes associated with brain development or function. Yet, since the RNA sequencing was performed on whole larvae rather than isolated brain samples, it is possible that our analysis missed genes expressed in low abundance in the brain. Nevertheless, our histological analyses revealed that loss of motile cilia-driven flow does not exert any major consequences on brain anatomy or cell proliferation. This contrasts with our investigation of zebrafish mutants lacking primary cilia after neurulation which revealed altered neurodevelopmental processes (43).

To understand whether motile cilia paralysis leads instead to physiological effects, we performed whole brain calcium imaging. We observed that the *smh* mutants exhibited reduced photic-evoked neuronal activity in the optic tectum and hindbrain regions. These brain regions are associated with visual processing and coordination (59, 60) respectively, and alterations in their neuronal responses could lead to impairments in sensory perception and motor coordination. Due to the body curvature of cilia mutants, experiments assessing motor coordination are unfortunately not possible. Interestingly, the reduced neuronal response was not confined to the neurons located in the midline in proximity to the ventricles, suggesting a more widespread impact of motile cilia-mediated flow. This raises intriguing questions about potential signaling mechanisms or inter-regional communication affected by altered cilia-mediated flow. Previous work in mice (13) showed that the migration of neuroblasts in the adult brain follows the flow of cerebrospinal fluid (CSF). They also reported that beating of ependymal cilia forms a concentration gradient which aids directional migration of neuroblasts. Hence, it is possible that altered neuronal migration could underly the physiological impact we observed. In our experiments, we investigated the retina and its projections to the brain to confirm that the reduced photic responses were not due to a defective retina. Notably, we found no defects in the retinal connections to the tectum in the motile cilia mutants. This analysis suggests that at least neuronal projections from the retina to the optic tectum were not affected upon loss of ciliary beating. In contrast to the retina, we observed disruption in axonal tracts projecting from the olfactory bulb to the habenula and randomized photic responses in the habenula of *smh* mutants as opposed to asymmetrical responses in controls. This raises the possibility that motile cilia are involved in axonal pathfinding in the habenula. Interestingly, we observed no asymmetry defects in the habenula of *elipsa* mutants, which loose cilia before motile cilia are formed in the brain, but after left-right asymmetry is established (43). This suggests that it is not motile cilia in the brain that regulate habenular asymmetry, but developmental processes downstream of the left-right organizer (46, 61-64). Notably, left-right organizer defects in *smh* mutants have been confirmed earlier (26). Future studies are needed to unravel how the left-right organizer regulates the establishment of habenular asymmetry (61-70). Given that the habenula plays a crucial role in various neurological processes, including sensory processing (38-40, 42, 65), circadian rhythms (71), mood regulation (72, 73), social interaction (74) and learning (75-77), the habenula may be an interesting brain region for further investigation in the context of cilia-mediated flow and brain asymmetry.

Our analysis of spontaneous activity showed no differences between control and mutant groups in any brain region hinting that cilia-mediated flow might not influence spontaneous activity in the brain. However, we did observe slightly increased correlation between cells in the telencephalon. Given that habenular circuits are closely interacting with telencephalic regions (38, 40), defects in habenular asymmetry may be linked to the slightly increased correlation between telencephalic neurons, which will require further investigations.

To further identify the mechanism underlying reduced neuronal activity, we investigated astroglial cells, which provide an essential support role in synapse formation and neurotransmission and thereby facilitate neural connectivity and communication within the nervous system (29, 78-81). We observed strikingly reduced astroglial activity in the *smh* mutant, which indicates potential disruptions in neuronal support, ion balance, and neurotransmitter regulation. In zebrafish, the majority of astroglia exhibit morphological traits reminiscent of mammalian radial glia, showcasing radial extensions that extend from the ventricular surface to the pial surface (29). Yet, increasing evidence support a role of zebrafish astroglia in neural circuit regulation (48, 49, 82-84), similarly to mammalian astrocytes. Notably, molecules such as norepinephrine (85-88), acetylcholine (89-91), and glutamate (92-94) have been proposed as primary triggers for activating astrocytes. Furthermore, activated astrocytes release gliotransmitters which in turn can modulate neural activity (95-97). The reduced astroglial activity observed in the *smh* mutant could potentially be linked to dysregulation of neurotransmitters, gliotransmitters or neuromodulators in the CSF (23, 98). The altered CSF flow caused by dysfunctional motile cilia may lead to changes in the availability and distribution of these signaling molecules near the ventricular walls, impacting astroglial responses and neural activity. In addition to molecules in the CSF, astroglia can also be activated through mechanosensors directly. Mechanical stimuli were shown to trigger calcium waves in glial cells (99). Additionally, work in mice revealed that the mechanosensor Piezo1 mediates Ca^2+^-dependent ATP release from astrocytes (54). Given that cilia mediated flow dominate near wall flow (4, 12, 15), this flow could exert mechanical forces on the ventricular walls thereby activating mechanosensors on glia cells and ultimately modulating neuronal activity. In the zebrafish, the primary cilia of astroglia and neuronal progenitors point toward the CSF-filled ventricles (8, 29, 100). Since cilia were shown to sense and respond to mechanical stimuli (101), one could hypothesize that primary cilia of astroglia are involved in this process. Taken together, we trust that our findings point to a crucial connection between motile cilia function and glial responses, warranting further investigation into the underlying mechanisms and identifying whether astroglia are regulated by the chemical nature of CSF or mechanical forces exerted by cilia mediate flow.

In conclusion, our comprehensive analysis of the *smh* mutant provides valuable insights into the consequences of impaired cilia-driven flow, particularly on neuronal activity and astroglial function. Understanding the impact of motile cilia dysfunction on brain physiology and neuronal responses, and its molecular underpinnings will contribute to further addressing neurodevelopmental disorders in ciliopathies in animal models and humans.

## Acknowledgements

We thank Valérie Wittamer for sharing the 4C4 antibody, our fish facility team for husbandry maintenance and technical support, and all members of the Jurisch-Yaksi and Yaksi laboratories for their feedback on this work and exchanging MATLAB codes. This work was supported by funding from an NTNU strategy grant (NJY) and The Research Council of Norway: RCN FRIPRO grant 314189 (NJY), RCN FRIPRO grant 314212 (EY).

## Author contributions

Conceptualization: NJY, PPD; Methodology: PPD, IJ, AN, AJ, EY; Formal analysis: PPD, IJ, AJ, NJY; Investigation: PPD, IJ, AN, NJY; Resources: NJY, EY; Data curation: PPD, IY, NJY; Writing - original draft: PPD, NJY; Writing - review & editing: all authors; Visualization: PPD, NJY; Supervision: NJY, EY; Funding acquisition: NJY, EY.

## Materials and Methods

Key Resource Table:

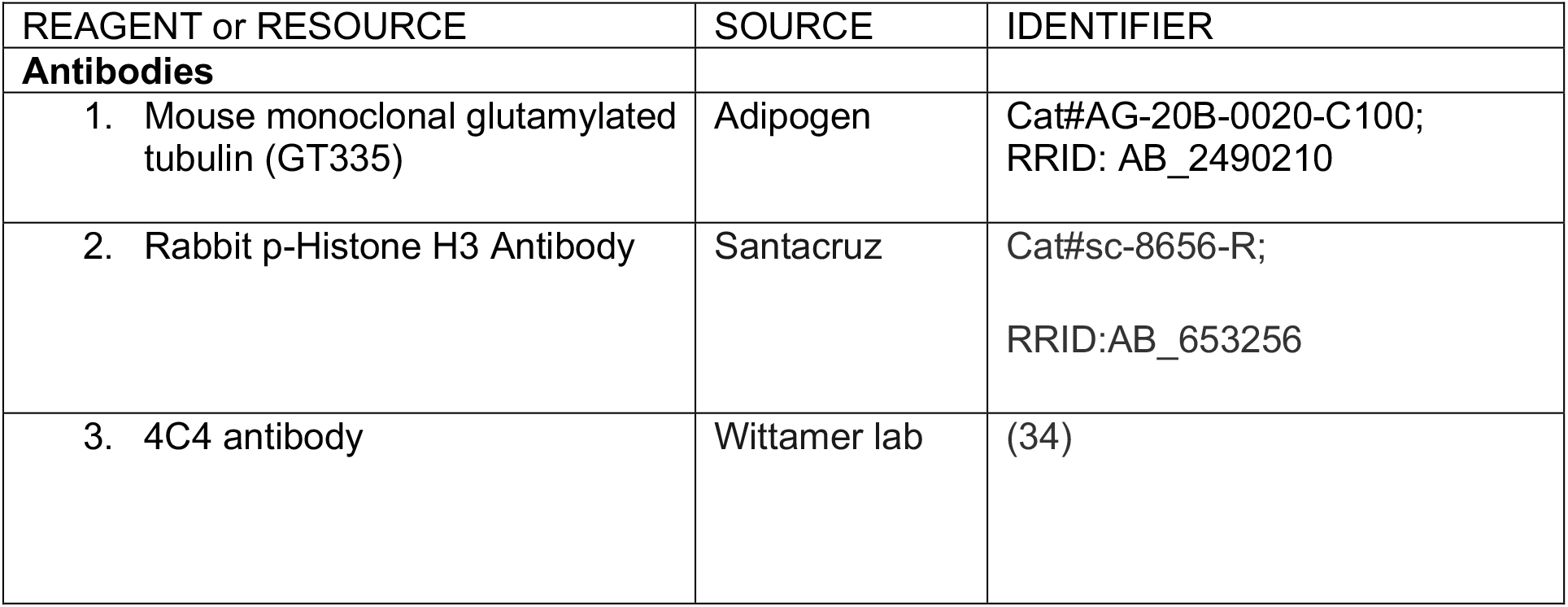
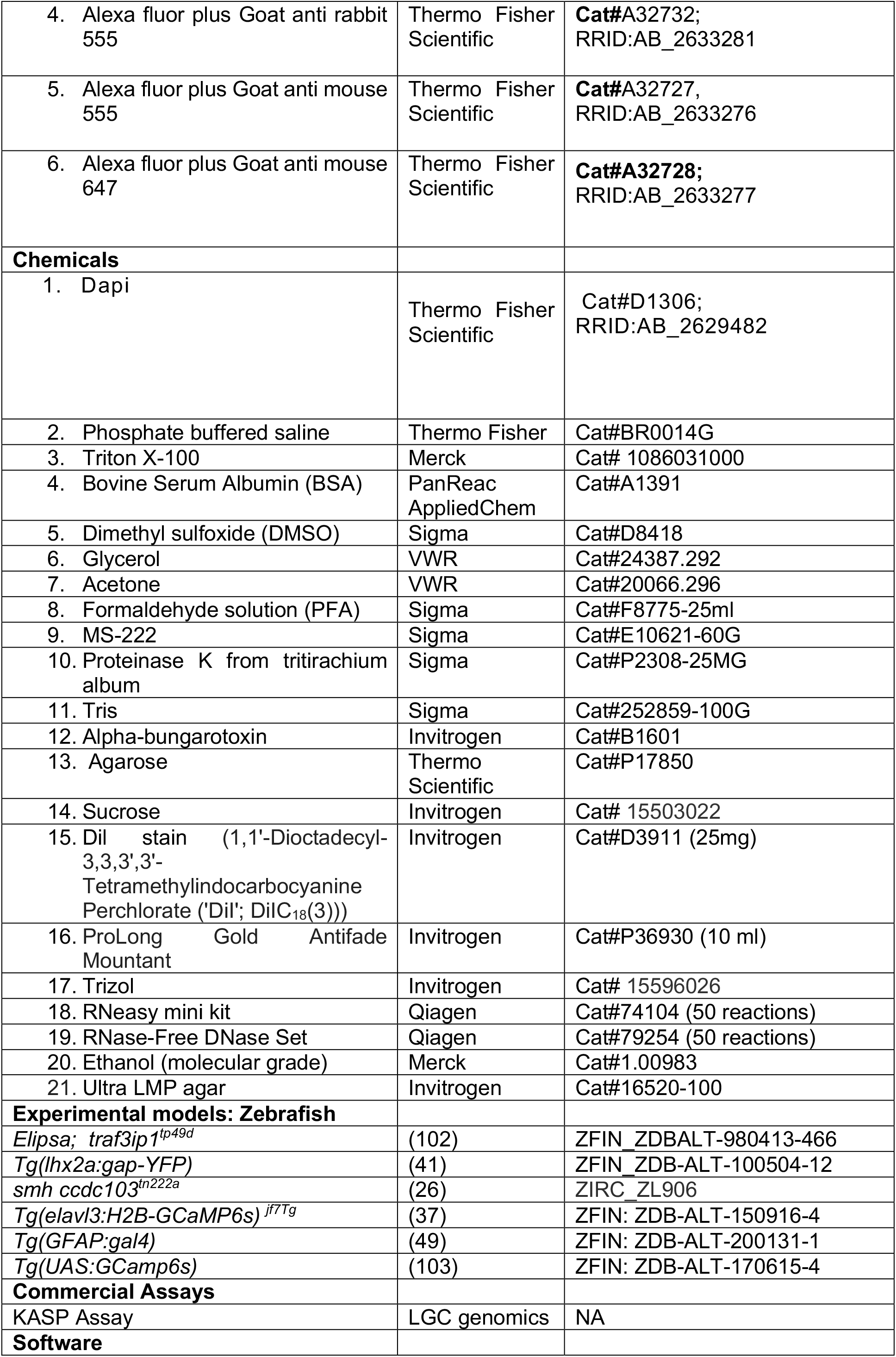
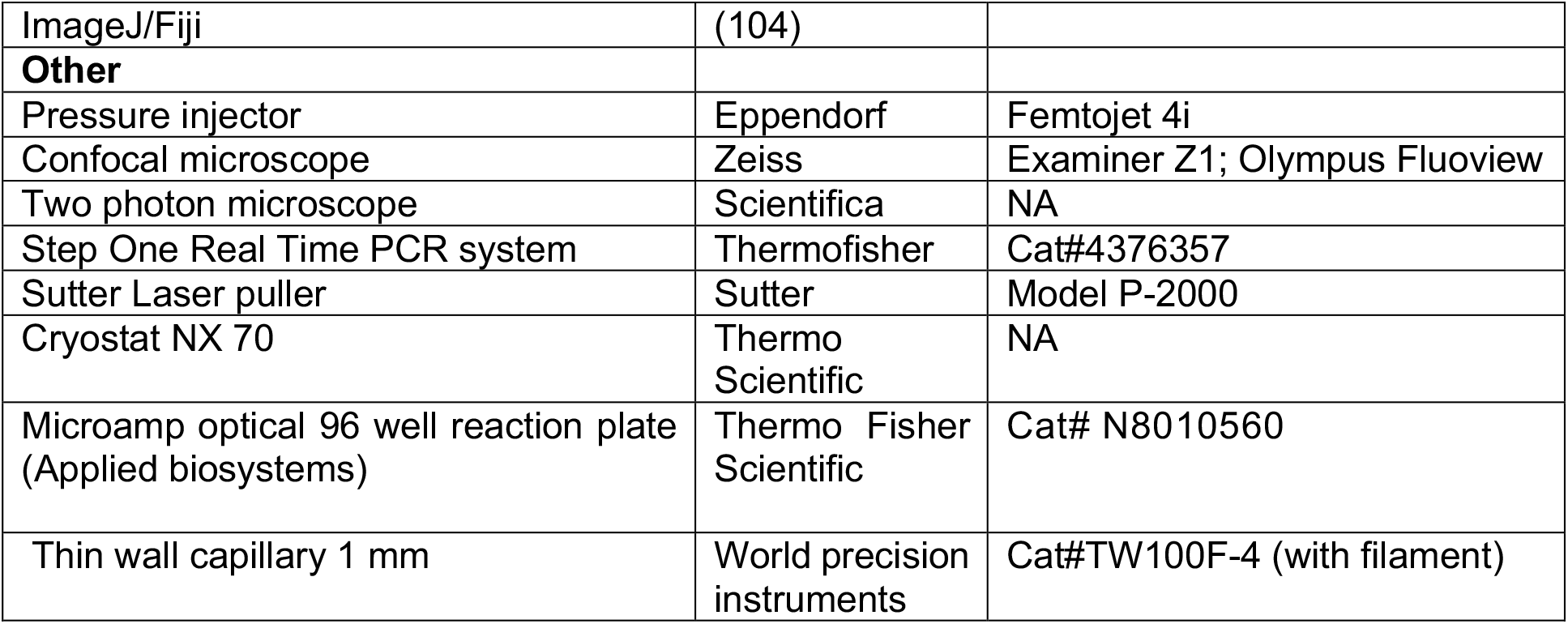

## Resource Availability

For inquiries about resources and reagents, please contact lead contact Nathalie Jurisch-Yaksi at.

## Methods

### Zebrafish Maintenance and Strains

The zebrafish, *Danio rerio*, were maintained in accordance with the guidelines set by the Norwegian Food Safety Authority (NFSA) and the European Communities Council Directive. The larval and adult zebrafish were raised under standard husbandry conditions at 28.5°C in 3.5 L tanks connected in a Techniplast Zebtech Multilinking system. The tanks were kept at constant pH 7 and 700 μSiemens with a 14:10 hr light/dark cycle to simulate optimal natural breeding conditions. From fertilization to 3 dpf, larvae were maintained in egg water (1.2 g marine salt and 0.1% methylene blue in 20 L RO water) and subsequently transferred to AFW (1.2 g marine salt in 20 L RO water). The fish lines used in the experiments were *smh ccdc103*^*tn222a*^ (received from Drummond lab, MGH), *Tg(elavl3:H2B-GcaMP6)*^*jf7Tg*^, *Tg(lhx2a:gap-YFP) and Tg(GFAP:gal4)Tg(UAS:GCamp6s)*. Two-photon calcium experiments were conducted with larvae obtained from in-crossing heterozygous *smh*^*+/-*^;*Tg(elavl3:H2B-GCaMP6s)* ^*jf7Tg*^ adult animals. For immunostaining and RNA seq, larvae were obtained by crossing heterozygous *smh*^*+/-*^ animals. Control groups consisted of either wild-type or *smh*^+/-^ fish obtained from the same breeding. For measuring glia activity *smh*^*+/-*^

;*Tg(GFAP:gal4)Tg(UAS:GCamp6s) fish line was used*.

### Genotyping

Genomic DNA (gDNA) was isolated from the samples using 100 μL of PCR lysis buffer (containing 1M tris pH-7-9, 0.5 M EDTA, tritonX-100, and Proteinase K 0.1mg/ml) overnight at 50°C. To halt the reaction, the samples were heated to 95°C for 10 minutes and then centrifuged at 13000 rpm for 2 minutes. The supernatant containing gDNA was utilized for KASP assays-based analysis. The samples were diluted (1:2) with water, and 3 μL of the diluted sample was used to perform the KASP assay following the manufacturer’s guidelines. The master mix for each sample well contained 5 μL of master mix, 0.14 μL of assay mix, and 1.86 μL of milliQ water.

### Antibody staining and confocal imaging

#### Immunostaining of the brain with cilia-specific antibodies

To perform immunostaining, zebrafish larvae were euthanized and fixed in a solution containing 4% paraformaldehyde solution (PFA), 1% DMSO, and 0.3% TritonX-100 in PBS (0.3% PBSTx) for 2 hours at room temperature or overnight at 4°C. After fixing, the larvae were washed with 0.3% PBSTx to remove any traces of the fixing solution. To permeabilize the samples, they were incubated for 10 minutes at -20°C with acetone. Subsequently, the samples were washed with 0.3% PBSTx (3×10 minutes) and blocked with 0.1% BSA in 0.3% PBSTx at room temperature. The samples were then incubated with primary antibodies overnight at 4°C. The antibodies used included mouse glutamylated tubulin (GT335, 1:400, for staining motile cilia), or rabbit p-Histone H3 Antibody (1:500, for staining dividing cells). On the following day, the samples were washed with 0.3% PBSTx (3×1 hour) and subsequently incubated with the secondary antibody (Alexa-labeled GAM488 plus or GAR555 plus Thermo Scientific, 1:1000) and 0.1% DAPI overnight at 4°C. After incubation with the secondary antibody, the larvae were washed (0.3% PBSTx, 3×1 hour) and transferred to a series of increasing glycerol concentrations (25%, 50%, and 75%) made in PBS. The stained larvae were stored in 75% glycerol at 4°C and imaged using a Zeiss Examiner Z1 confocal microscope with a 20x plan NA 0.8 objective. Multiple images were stitched using Fiji software (105). For a detailed protocol, please refer to (106).

#### Quantification of pH3+ and microglial cells and measuring brain morphology

After image acquisition, the pH3+ and microglial cells were counted using the cell counter function in Fiji (https://imagej.nih.gov/ij/plugins/cell-counter.html). Brain morphology measurements were performed on Z-stack images using the straight-line tool in Fiji.

#### Cryo-sectioning and staining of the retina

The protocol for cryo-sectioning was adapted from (107). Zebrafish larvae were euthanized using ice water (4°C) and then fixed in 4% PFA in PBS overnight at 4°C. After fixing, the samples were washed three times for 1 hour each with PBS. The fish were embedded for cryoprotection using a solution of 1.5% agarose and 5% sucrose in RO water. The larvae were transferred to a cryomold, and the embedding solution was added. After cooling, the mold was cut into tiny blocks, and the blocks were transferred to a storage solution (30% sucrose in PBS) and stored overnight at 4°C until the samples sank to the bottom of the tube. The next day, the blocks containing the samples were snap-frozen with liquid nitrogen and stored at -20°C until further use. Sections of 10 µm thickness were cut at -30°C blade and chamber temperature using a cryostat. The sections were placed on super frost slides and stored at -20°C until further use. The cryosection slides were thawed at room temperature for 30 minutes. The slides were washed with PBS without detergent four times for 5 minutes each to remove any traces of freezing medium. The outer segments were stained using DiI (5mg/ml diluted 1:100-1:200 in PBS) stain for 20 minutes at room temperature, followed by three washes with 1X PBS for 5 minutes each. The slides were then dried and mounted with Prolong Gold. The slides were left overnight to dry before imaging.

#### RNA Sequencing and transcriptomic analysis

To isolate RNA for sequencing, 30 larvae at 4 dpf stage were collected in a 1.5ml tube and kept on ice. To lyse the samples, 500μL trizol was added, following which the samples were homogenized through a 27-gauge needle until the mixture looked uniform. After adding another 500μL trizol, the samples were incubated for 5 minutes at room temperature. The samples were then treated with 200μL chloroform, and the tube was rocked for 15 seconds to mix the contents. The tubes were incubated for 2 minutes at room temperature and then centrifuged for 15 minutes at 12000rpm at 4°C. After centrifugation, the upper aqueous phase containing RNA was mixed with equal amounts of 100% ethanol and loaded onto an RNA spin column (Qiagen). The spin column was further treated to remove any DNA contamination by washing with RW1 buffer and DNase enzyme in RDD buffer. After incubation, the samples were treated with RPE buffer and centrifuged to remove any residual buffer left in the column. RNA was extracted from the column using nuclease-free water. The concentration and quality of the extracted RNA were determined using Nanodrop and bioanalyzer, respectively. The samples were then sequenced using BGI’s DNBSEQTM Technology using the Dr Tom data visualization and analysis platform provided by BGI. The filtering of sequencing data was done using SOAPnuke, v1.5.2, using the parameters Parameters -l 15 -q 0.2 -n 0.05. The Hierarchical Indexing for Spliced Alignment of Transcripts software (HISAT2 v2.0.4 with parameters:--sensitive --no-discordant --no-mixed -I 1 -X 1000 -p 8 --rna-strandness RF) was used for mapping RNA-seq reads. We used Bowtie2 (108) (Version:v2.2.5, Parameters:-q --sensitive --dpad 0 --gbar 99999999 --mp 1,1 --np 1 --score-min L,0,-0.1 -p 16 -k 200) to map the clean reads to the reference gene sequence (transcriptome), and then RSEM (109) (Version:v1.2.8, Parameters:-p 8 --forward-prob 0 --paired-end) to calculate the gene expression level of each sample. The count matrix included genes that were selected based on their expression of average count per million of more than 1 across all samples. The resulting matrix was normalized, and log transformed using the voom algorithm from the limma package of Bioconductor (110, 111). For performing gene ontology (GO) analysis, the differentially expressed genes having a p-adjusted value of 0.1 was selected and analyzed using the shiny GO gene set enrichment tool (http://bioinformatics.sdstate.edu/go/) (112).

#### Two-photon calcium imaging

Two-photon calcium imaging was performed on *smh*^*+/-*^;*Tg(elavl3:H2B-GCaMP6s)* and *smh*^*+/-*^

;*Tg(GFAP:Gal4);Tg(UAS:GCaMP6s)* 4 dpf zebrafish larvae. The larvae were paralyzed upon injection of α-bungarotoxin (Invitrogen BI601, 1 mg/ml) and embedded in 1.5% low melting point agarose in mounting chambers (Fluorodish, World Precision Instruments) using a plastic microcapillary tip, as described in previous work (113). After agarose solidification, AFW was added on top. The mounting chamber was allowed to stabilize for 15 minutes under the two-photon microscope before recording. The recordings were performed in a two-photon microscope (Scientifica) using a 16× water immersion objective (Nikon, numerical aperture 0.8, Long Working Distance 3.0, plan) and a Ti:Sapphire laser (MaiTai Spectra-Physics) tuned at 920 nm. Multiple images were acquired at a frame rate of 30.85 frames per second and a volume rate of 3.86 Hz. The total recording time was 40 minutes (74000 frames). Ongoing activity was recorded for 10 minutes in darkness, followed by five stimuli each of 60 seconds using a red light-emitting diode light (LZ1-00R105, LedEngin; 625 nm) at minutes 10, 15, 20, 25, and 30 respectively as described in (36). Animals without cerebral blood flow after the experiments were excluded from the analysis.

### Data analysis

Two-photon microscopy images were aligned using a previously reported algorithm (40, 113). The recordings were then screened manually to check for movement and Z drift. Unstable recordings were discarded from the analysis. All analyses were performed on MATLAB. Neurons were segmented using a pattern recognition algorithm adapted from (114) with torus or ring shaped neuronal templates (42). The identified neurons were then set apart into different brain regions as described in (38). Clustering of neurons was performed using k-means clustering algorithm based on their activity (42). For detecting light responses, we used the 5 second of fluorescence preceding the stimuli as baseline and average of the 5 stimuli for each neuron. The 5s average baseline was used to calculate dF/F. Responsive cells were selected based on their response during the 10s following the light onset or offset. Cells were classified as responsive if their average activity during the 10s window was greater than 2 ^*^ standard deviation of the baseline. Neurons with an average activity below 2 ^*^ standard deviation of the baseline were considered as inhibited.

For measuring spontaneous activity, we excluded the first two minutes of the recordings (to avoid artefacts from the microscope laser turning on). We selected frames 460-2304 (a total of 7.5 minutes of ongoing activity). dF/F was calculated using as baseline the 8^th^ percentile of a moving window as previously described (40, 115).The data was resampled to 1fps using the resample function in MATLAB. Calculations for the activity of cells were done on resampled data. A cell was considered active if it had an activity higher than a threshold of 4 ^*^ baseline. We then calculated the percentage of cells that are highly active (active more than 50% of the time) or inactive (active less than 10% of the time) per brain region. The correlation versus distance between neurons was calculated up to 60μm per brain region **(figure S2)**.

### Data analysis for glia activity

For two photon recordings measuring astroglial activity, the same light stimulation protocol was used as mentioned above. ROIs were drawn around the telencephalon, thalamus, optic tectum, and hindbrain to extract fluorescence specifically from the soma of the glia cells lining the ventricles of the brain **(figure 4A)**. The activity was averaged across all ROIs.

### Electroretinography

Electroretinography (ERG) experiments were conducted on both control and *smh* mutant larvae at the 4dpf stage. Initially, healthy control and *smh* mutant larvae were selected and anesthetized using MS222. Subsequently, the anesthetized larvae were placed on a wet filter paper (VWR, 516-0848) situated on a FluoroDish (VWR, FD35PDL-100). To secure them in place, their trunks were covered with paper towels. A silver-coated wire, previously treated with sodium hypochlorite for 10 minutes, was used as a reference electrode and positioned in the same bath as the larvae. Recording electrodes were pulled from glass capillaries (WPI, TW100F-4) with an opening size of 15-30 μm and were filled with artificial fish water (AFW) containing MS222. These electrodes were then placed on the cornea of the eye using a motorized micromanipulator from Scientifica. Before recording, the larvae were allowed to dark-adapt for over 10 minutes. Voltage measurements were obtained using a MultiClamp 700B device from Molecular Devices, featuring a 2 kHz low-pass filter and digitized at 10 kHz. A series of light stimuli were initiated at the 5th second of the recording. These stimuli were generated by a blue LED using a pulse generator (Master-8, AMPI). Each stimulus, with a light intensity ranging from 0.005 to 0.007 mW, lasted for 1 second, with 5-second intervals between stimuli. To ensure accuracy, at least 5 recordings were performed for each larva, and the trial exhibiting the most representative responses was selected for further analysis. Custom MATLAB scripts were employed for data acquisition and subsequent analyses. To quantify the voltage responses, a 200-millisecond baseline before the light stimulus was utilized to normalize the traces. The responses to 3 light stimuli were averaged. The average voltage responses for both control and mutant larvae were calculated within the 0 to 200-millisecond timeframe after the onset of the light stimuli, which primarily represents the strong b-wave indicative of the ON bipolar cell response.

### Quantification and statistical analysis

Statistical analysis was done using MATLAB. Wilcoxon rank-sum test was used for nonpaired analysis. Probability of *p* < 0.05 was considered statistically significant.

