## Supplementary figures for "Cilia-mediated cerebrospinal fluid flow modulates neuronal and astroglial activity in the zebrafish larval brain"

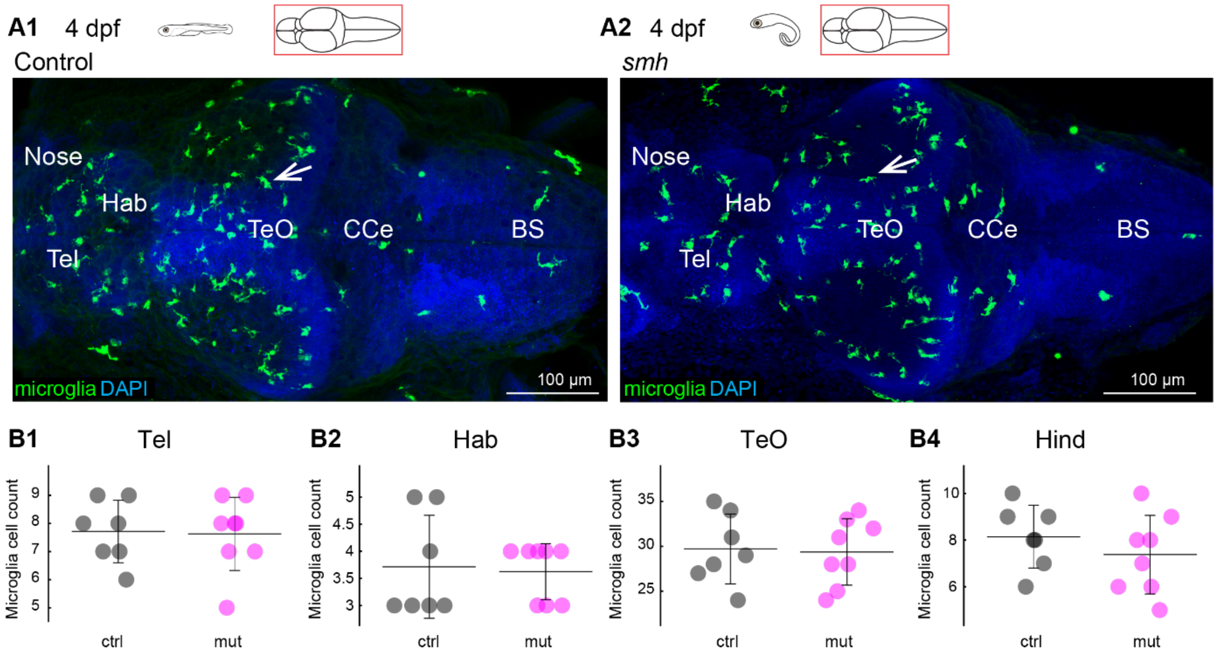


**Supplementary figure 1: Loss of cilia motility does not impact microglia in the brain.**

**(A1, A2)** Staining for microglia cells indicated by arrows in **A1, A2** using the 4C4 antibody. **(B1-B4)** Cell count for microglia cells in telencephalon **(B1)**, habenula **(B2)**, optic tectum **(B3)** and hindbrain **(B4)**. *n* =7 controls, 8 mutants. Mean +/- standard deviation is indicated on scatter plots. Tel, Telencephalon; Hab, Habenula; Teo, Optic Tectum; CCe, Corpus Cerebelli; BS, Brain stem.

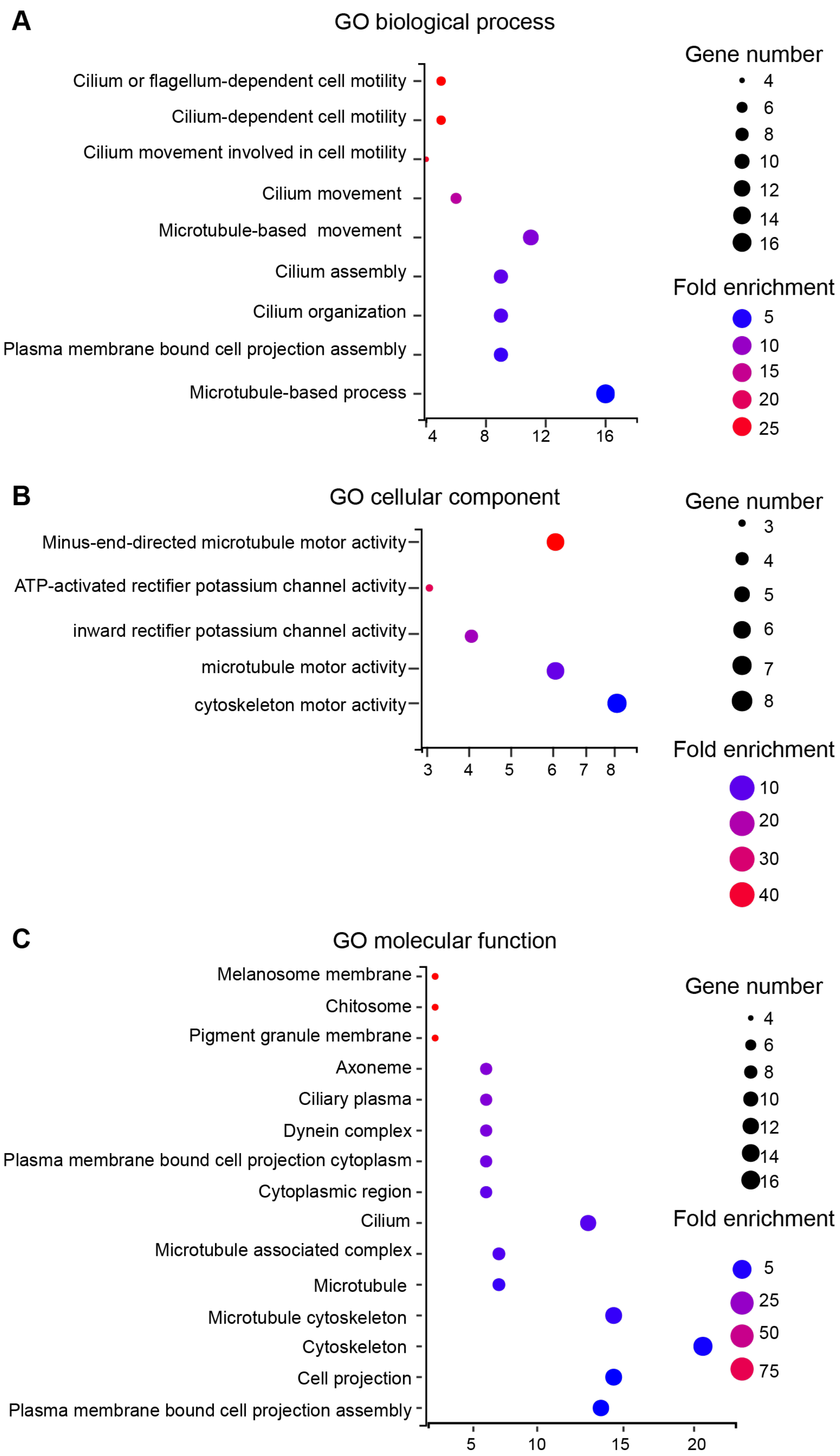
A
GO biological process
Cilium or flagellum-dependent cell motility
Cilium-dependent cell motility
Cilium movement involved in cell motility
Cilium movement
Microtubule-based movement
Cilium assembly
Cilium organization
Plasma membrane bound cell projection assembly
Microtubule-based process
4
8
12
16
Gene number
4
6
8
10
12
14
16
Fold enrichment
5
10
15
20
25
B
GO cellular component
Minus-end-directed microtubule motor activity
ATP-activated rectifier potassium channel activity
inward rectifier potassium channel activity
microtubule motor activity
cytoskeleton motor activity
3
4
5
6
7
8
Gene number
3
4
5
6
7
8
Fold enrichment
10
20
30
40
C
GO molecular function
Melanosome membrane
Chitosome
Pigment granule membrane
Axoneme
Ciliary plasma
Dynein complex
Plasma membrane bound cell projection cytoplasm
Cytoplasmic region
Cilium
Microtubule associated complex
Microtubule
Microtubule cytoskeleton
Cytoskeleton
Cell projection
Plasma membrane bound cell projection assembly
5
10
15
20
Gene number
4
6
8
10
12
14
16
Fold enrichment
5
25
50
75

**Supplementary figure 2: Gene Ontology (GO) analysis reveals differentially regulated genes involved in ciliary movement and dynein activity.**

**(A)** Top 20 gene Ontology (GO) biological process **(B)** Top 20 GO cellular component **(C)** Top 20 GO molecular function differentially regulated in the *smh* mutant.

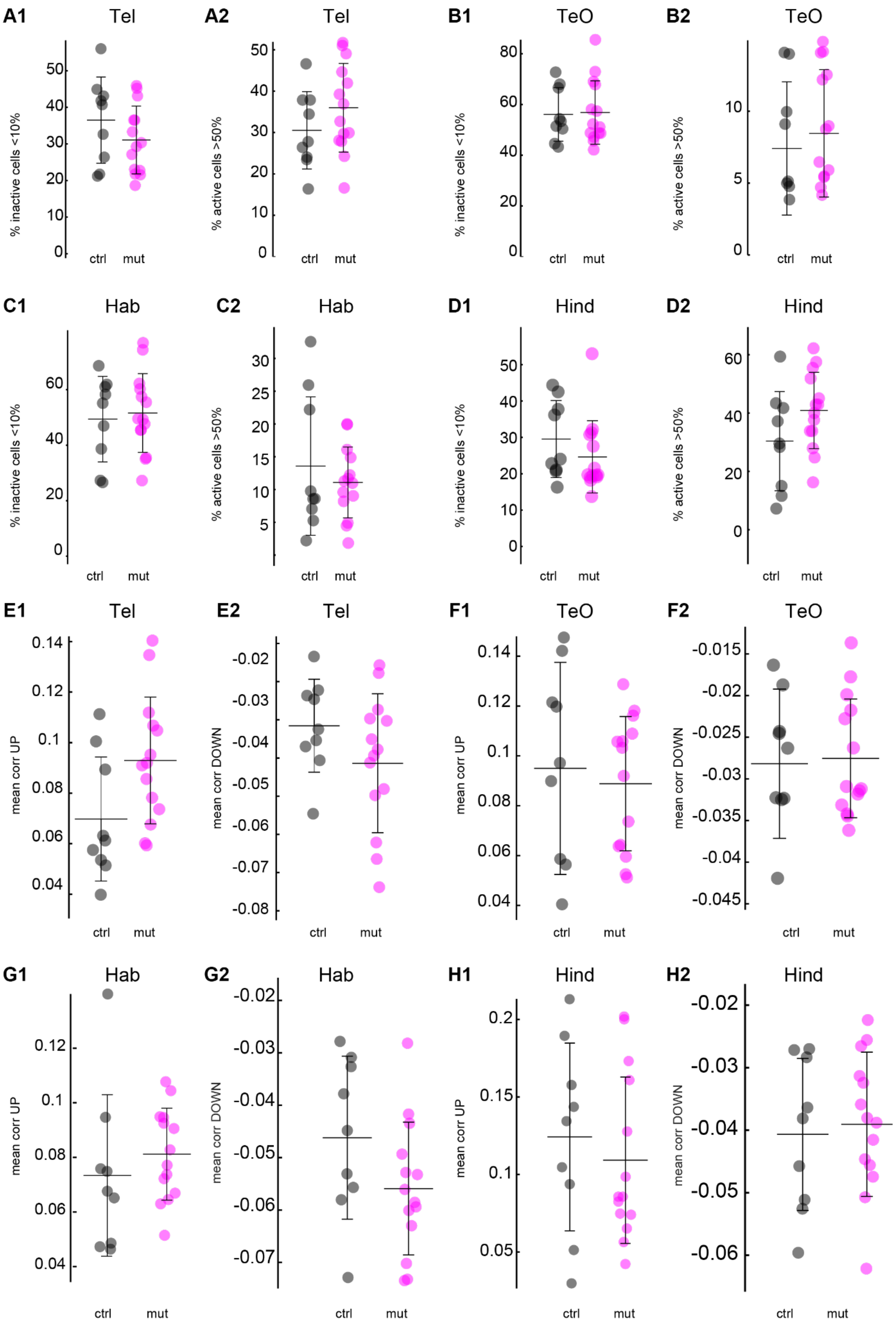
A1
Tel
% inactive cells <10%
0
10
20
30
40
50
ctrl
mut
A2
Tel
% active cells >50%
0
10
20
30
40
50
ctrl
mut
B1
TeO
% inactive cells <10%
0
20
40
60
80
ctrl
mut
B2
TeO
% active cells >50%
0
5
10
ctrl
mut
C1
Hab
% inactive cells <10%
0
20
40
60
ctrl
mut
C2
Hab
% active cells >50%
5
10
15
20
25
30
ctrl
mut
D1
Hind
% inactive cells <10%
0
10
20
30
40
50
ctrl
mut
D2
Hind
% active cells >50%
0
20
40
60
ctrl
mut
E1
Tel
mean corr UP
0.04
0.06
0.08
0.1
0.12
0.14
ctrl
mut
E2
Tel
mean corr DOWN
-0.08
-0.07
-0.06
-0.05
-0.04
-0.03
-0.02
ctrl
mut
F1
TeO
mean corr UP
0.04
0.06
0.08
0.1
0.12
0.14
ctrl
mut
F2
TeO
mean corr DOWN
-0.045
-0.04
-0.035
-0.03
-0.025
-0.02
-0.015
ctrl
mut
G1
Hab
mean corr UP
0.04
0.06
0.08
0.1
0.12
ctrl
mut
G2
Hab
mean corr DOWN
-0.07
-0.06
-0.05
-0.04
-0.03
-0.02
ctrl
mut
H1
Hind
mean corr UP
0.05
0.1
0.15
0.2
ctrl
mut
H2
Hind
mean corr DOWN
-0.06
-0.05
-0.04
-0.03
-0.02
ctrl
mut

**Supplementary figure 3: No differences in spontaneous activity and correlation between neighboring neurons in the brain of motile cilia mutants.**

(**A1-D2)** Quantification of inactive and active cells for the **(A1-A2)** telencephalon, **(B1-B2)** optic tectum and **(C1-C2)** habenula and **(D1-D2)** hindbrain regions, controls in black and *smh* mutant in magenta. *n* = 9 controls and 14 mutants. **(E1-H2)** Quantification of mean positive (UP) and negative (DOWN) correlation for cells located within 60 µm distance in the **(E1-E2)** telencephalon, **(F1-F2)** optic tectum, **(G1-G2)** habenula and **(H1-H2)** hindbrain regions, controls in black and *smh* mutant in cyan. *n* = 9 controls and 14 mutants. Mean +/- standard deviation is indicated on scatter.

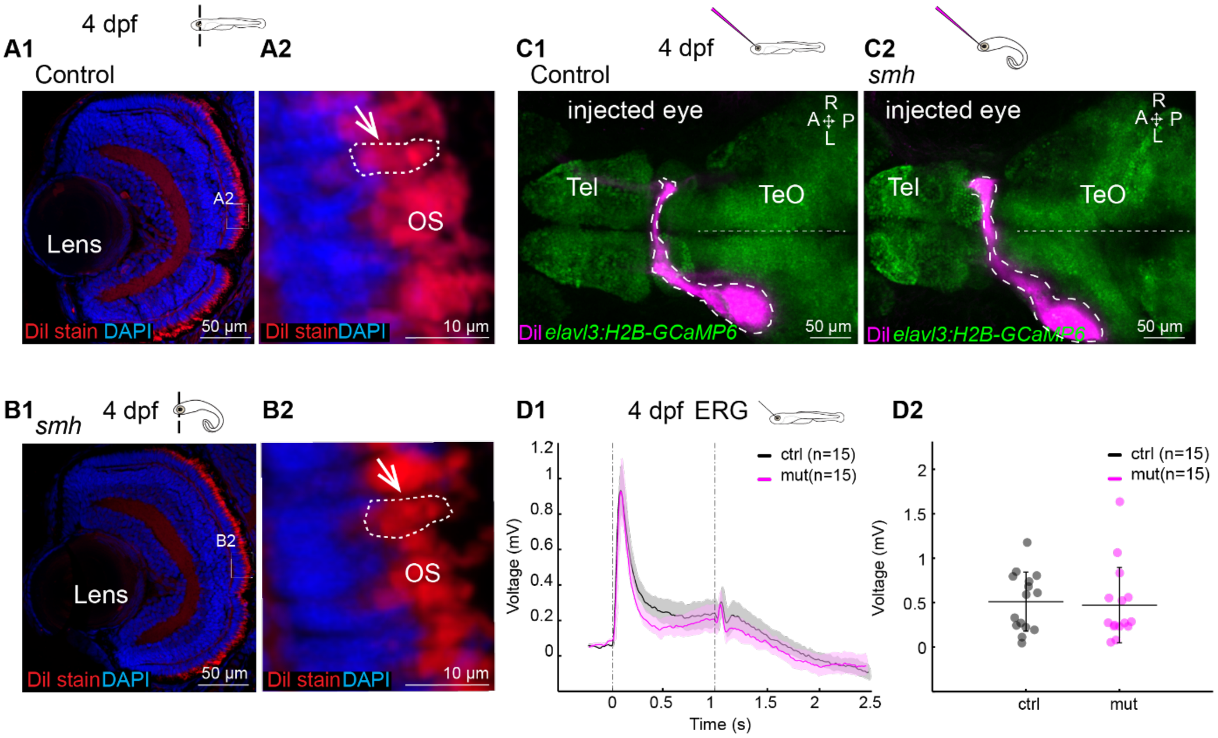


**Supplementary figure 4: *smh* mutant display no morphological defects of the retina outer segments and a normal ERG response.**

**(A1-B2)** DiI staining of 4 dpf retina cryosection to stain the outer segments. Section through the whole retina or the photoreceptor layer of a representative control **(A1-A2)** and *smh* mutant **(B1-B2)**. Normal morphology of outer segments (OS) is indicated using arrow and dashed lines. *n* = 8 controls and 10 mutants. **(C1-C2)** DiI injection into the eye at 4 dpf to stain the axonal connections (represented in dotted lines) which cross the midline and innervate the contralateral optic tectum. Here is shown a representative control **(C1)** and *smh* mutant **(C2).** *n* = 11 controls and 12 mutants. **(D1-D2)** Electroretinography (ERG) recordings in a 4 dpf retina. **D1** Average response of electrical activity (+/- standard error of the mean as the shaded region) to 1 second light stimulation for control (black) and mutant (magenta). **D2** Average electrical responses for the 200 msec following the light ON stimulus for all control fish (black) and *smh* mutant (magenta) showing normal activity in the mutant. Mean +/- standard deviation is indicated on scatter. *n* =15 controls and 15 mutants. Tel, Telencephalon; Teo, Optic Tectum.

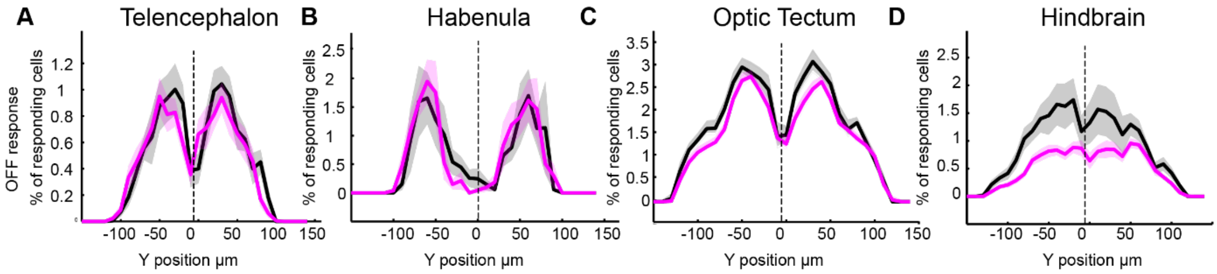


**Supplementary figure 5: Decreased neural activity in the optic tectum and hindbrain regions extends beyond the cells lining the ventricles.**

Y position for responding cells in the OFF response for telencephalon **(A1)**, Habenula **(A2)**, optic tectum **(A3)**, and hindbrain **(A4).** Dotted line at 0 represents the position of the cells lining the ventricles.

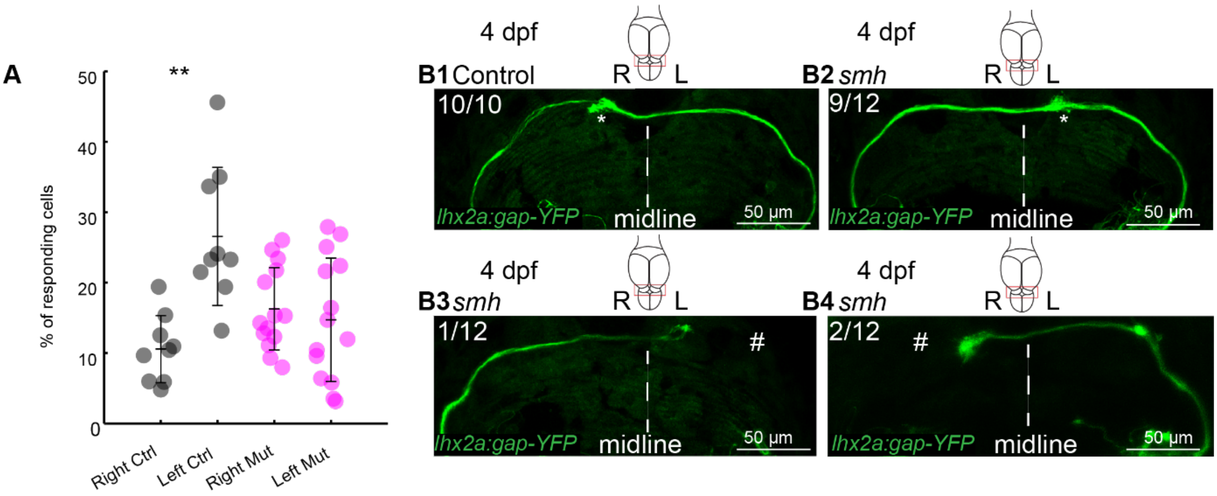


**Supplementary figure 6: The habenular response to light and axonal projections from the olfactory bulb to the habenula exhibits reversed orientation.**

**A** Percentage of cells that are activated during ON response in either left or right habenula. **: $p$ < 0.01 according to Wilcoxon Rank Sum test. Mean +/- standard deviation is indicated on scatter plots. **B1-B4** *Lhx2a:gap-YFP* line which labels axons from the olfactory bulb to the habenula in control **(B1)** and *smh* background **(B2-B4)**. Numbers of observations is indicated in the upper left corner of the images. *: axonal projections in the habenula, #: axonal projections to habenula from the olfactory bulb, R: Right, L: left.

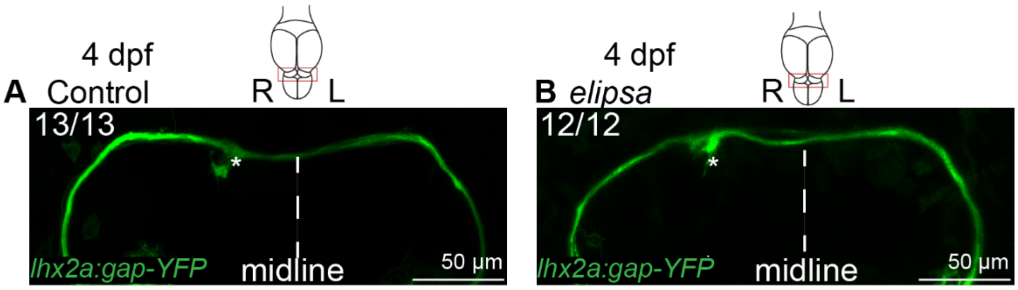


**Supplementary figure 7: The asymmetrical axonal projections from the olfactory bulb to the habenula are maintained in the *elipsa* mutant.**

*Lhx2a:gap:YFP* line which labels axons from the olfactory bulb to the habenula in control **(A)** and *elipsa* background **(B**). *: axonal projections in the habenula, R: Right, L: left.

**Supplementary Table 1:** Transcriptomic analysis of differentially regulated genes between control and *smh* mutant larvae.
